# The TREM-receptor family interactome in the human brain

**DOI:** 10.64898/2026.10.28.762136

**Authors:** Kim M.A. de Kleijn, Saskia A. Weller, Remco V. Klaassen, Frank T.W. Koopmans, August B. Smit, Ruud H. Wijdeven

## Abstract

The Triggering Receptors Expressed on Myeloid cells (TREMs) are a family of cell-surface receptors instrumental in the activation of different myeloid cells. TREM2, its most studied member, is genetically linked to Alzheimer’s disease (AD), and its interactions with Amyloid-β and APOE have placed it at the forefront of microglia dysfunction in the brain. However, the ligands and binding partners of the other TREM members (TREM1, TREML1, TREML2 and TREML4) are less well defined. Here, we used human AD and non-AD brain lysate to profile the interactome of the ligand binding domains of the different TREM receptors. Besides identifying known ligands for TREM1 (PGLYRP1), TREM2 (C1QL3, LGALSL) and TREML1 (Fibrinogens), we identified complement components and the apolipoproteins APOA1 and APOA4 as novel interactors of TREML1, and heparan sulfate modified proteins as binding partners for TREML2. Furthermore, similar to TREM2, all the TREMLs interacted with the apolipoproteins APOE and Clusterin. Using *in vitro* pulldowns with heparin, we confirmed that specifically TREML2 interacted with heparan sulfates, in a manner unaffected by the AD-linked S144G mutation. Together, our proteomics approach identified novel interactors for TREML1 and TREML2, cementing the role of the different TREM-proteins in lipid trafficking and immunity.

## Introduction

Activation and modulation of innate immune cells like microglia is governed by their numerous cell-surface receptors interacting with different ligands. An important class of these receptors is the TREM receptor family, in humans consisting of TREM1, TREM2, TREML1, TREML2 and TREML4. These receptors are expressed on different types of myeloid cells such as neutrophils, monocytes, and microglia, and fine-tune their activity ^1, 2^. All harbor a single immunoglobulin-like domain important for ligand binding, and most (except TREML1 and TREML2) contain an immunoreceptor tyrosine-based activation motif (ITAM) to interact with DAP12 (TYROBP) and drive intracellular signaling cascades ^3–5^. Dysfunction or activation of several TREM-receptors has been linked to brain disease ^6–8^. TREM2 is highly expressed in microglia and TREM1 is strongly upregulated in response to inflammation, while the TREM-like receptors are expressed at low levels in microglial subtypes.

Among the TREM family, TREM2 is the most well studied, due to its function in disease-associated microglia (DAM) often found in neurodegenerative conditions such as Alzheimer’s disease (AD) ^9, 10^. Several genetic variants in TREM2 that affect ligand binding and activation are linked to increased AD risk ^7, 11, 12^, showcasing its important role in AD development. TREM2 interacts with several different ligands, including lipid transporters (APOE, APOJ/Clusterin), amyloid-β, phospholipids, LGALS3 and complement factor C1q ^13–17^. Functionally, by binding to these different ligands, TREM2 is involved in lipid homeostasis, amyloid-β uptake, and the pruning of synapses ^18^.

In contrast to TREM2, TREM1 ligand engagement promotes (neuro)inflammation by boosting the transcriptional programs orchestrated by classical pro-inflammatory stimuli such as IL-1β and lipopolysaccharides (LPS) ^19^. TREM1 is expressed primarily in neutrophils, monocytes and macrophages, but pro-inflammatory stimulation upregulates its expression in many cell types, including microglia. Its only known endogenous ligand is Peptidoglycan Recognition Protein 1 (PGLYRP1), a pattern recognition molecule with antibacterial properties. The naturally existing affinity of PGLYRP1 for TREM1 is increased upon binding of bacterial peptidoglycans, making it a sensor for bacterial infection. In the brain, TREM1 has been shown to regulate neuroinflammation and ageing ^6, 19^, and its expression is upregulated during AD progression ^8^, in agreement with the increased microglia activation observed during the process.

The functions of the other part of the family, the TREM-like receptors, are less well characterized. TREML1 is primarily expressed on blood platelets, and functions to stimulate platelet aggregation and blood clotting by interacting with fibrinogens ^20, 21^. In AD, a genetic variant that increases TREML1 expression has been linked to a decreased risk in developing AD, but this variant also increases TREM2 expression, so it is unclear which TREM is responsible for this effect ^22^. Another TREM-like, TREML2, has a broader expression in different types of myeloid cells, and has recently been shown to promote pro-inflammatory signaling, in part by cooperating with Toll Like Receptor (TLR)-mediated signaling ^23^. Depletion of TREML2 decreased pathology in an AD mouse model ^24^, and a missense variant in the TREML2 gene (S144G) is protective for AD ^25, 26^. However, no direct ligands for the receptor have been established, so it is unclear what molecular mechanism drives TREML2 activation. Similarly, no ligand has been identified for the TREM-like protein TREML4, which was identified in two different genetic screens as important for pro-inflammatory TLR-signaling and sepsis ^27, 28^, but a mechanistic basis for this is lacking.

In case of TREM1 and TREM2, ligand binding leads to phosphorylation of DAP12 and subsequent downstream signaling, with signal termination orchestrated by TREM2 internalization or cleavage of the extracellular domain by metalloproteases (ADAM10 and ADAM17) ^29^. The latter leads to the presence of a soluble TREM (sTREM), and for sTREM2 its levels in the CSF of AD patients were predictive of disease state ^30^. Others have shown that sTREM2 turns on NFκB-signaling in microglia and ameliorates AD pathology ^31, 32^, suggesting that it functions as a signaling molecule to a yet unknown receptor. Also sTREM1 regulates microglia function, via PI3K-AKT signaling ^33^, again via an unknown receptor. The other TREM-receptors are potentially also cleaved upon ligand binding, so sTREML1, sTREML2 and sTREML4 could also influence inflammation via yet unidentified receptors.

Given the importance of the TREM receptors in (neuro)inflammation and AD, we ought to characterize their interactomes in the brain. To do so, we produced recombinant TREM extracellular domains (ECD)s in human cells, and used these to pull down interacting proteins from human brain tissue. This method allowed us to identify both transmembrane and soluble interactors of the (s)TREM proteins. Besides known interactors, we identified the complement system and APOA1/APOA4 proteins as novel interactors for TREML1, and identified heparan sulfate modified proteins as an important class of interactors for TREML2. Together, these findings solidify a broader role for the TREM proteins in lipid trafficking and (innate) immune activation.

## Methods

### Brain lysate preparation

Human postmortem brain tissues were derived from the Netherlands Brain Bank and stem from 2 AD and 2 non-AD subjects, from two brain regions (hippocampus and striatum, both sexes and various Braak stages; Table 1). 250 mg of each of the 4 brain tissues was homogenized in 5 ml NP-40 lysis buffer (1% NP-40 (nonionic polyoxyethylene surfactant), 50 mM Tris-HCl, pH 8.0, 150 mM NaCl, protease and phosphatase inhibitors) using an automated douncing machine to ensure maximal and consistent lysis across the tissues (12 strokes at 900 rpm). The homogenates were incubated for 1h at 4 °C, centrifuged at 18,000 g at 4 °C for 15 min, and total protein concentrations were measured via a BCA assay according to the manufacturer’s instructions (Thermo Fisher Scientific). Concentrations were adjusted to 1 mg/ml and all four brain lysate supernatants were pooled for the pulldowns.

**Table 1.** Characteristics of the human brain tissue blocks used for protein lysate preparation.

| Sample | Sex | Age | Braak stage | APOE status | Diagnosis | Brain region |
| --- | --- | --- | --- | --- | --- | --- |
| S99/086 | F | 82 | 5 | 44 | Alzheimer's disease | Striatum |
| S15/027 | F | 76 | 2 | 33 | Non-demented control | Caudate nucleus |
| S21/051 | M | 77 | 1 | 33 | Non-demented control | Hippocampus |
| S22/002 | M | 89 | 3 |  | Alzheimer's disease | Hippocampus |

### TREM constructs

The extracellular domains of the different TREM receptors were cloned into a mammalian AVI-6xHIS vector, which was made by swapping the GFP in the GFP-N1 vector for a linker and AVI-6xHIS sequence (GSTGGSLHHILDAQKMLWNHRDSGSGHHHHHH). For all TREMs except TREM2, the extracellular domain was defined using the uniprot annotation, while for TREM2 the known, cleaved extracellular domain sequence was used. TREM1 (aa 1-205), TREM2 (aa 1-157) and TREML1 (aa 1-162) were cloned directly before the AVI-6xHIS sequence. For TREML2 (aa 19-209) and TREML4 (aa 25-183), the signal sequence of HA (MKTIIALSYIFCLVFA) was used instead of the native signal sequence because secretion using the endogenous signal sequence was inefficient. Secreted BirA-Flag was a gift from Gavin Wright (Addgene plasmid # 64395), and the Flag-C1 control vector was described previously ^34^.

### TREM protein production and pulldown

HEK293T cells (maintained in DMEM medium with 8% fetal calf serum) were co-transfected with the secreted BirA construct and each individual TREM-construct using polyethelineimine (PEI) at a 1:3 DNA:PEI ratio. The day after transfection, medium was replaced by Opti-MEM with 50 µM biotin to promote TREM biotinylation. Medium with secreted TREM-proteins was harvested, filtered through 0.22 µm syringe filters to remove cells and debris, and then purified over 10 kDa spin filters to remove free biotin from the medium. The concentration of the biotinylated TREMs in the HEK293T medium was measured via western blot and detection with a Streptavidin-aR800 dye (Li-COR biosciences).

Per pulldown, 20 ul Neutravidin beads (29201; Thermo Fisher Scientific), 1 ug freshly prepared TREM-protein and 1 mg freshly prepared cleared brain lysate was used. Neutravidin beads were first washed twice with 0.5% NP-40 wash buffer (150mM NaCl, 0.5% NP-40, 5mM MgCl2, 50mM Tris pH 8) and then blocked with 0.5 ml freshly prepared chicken egg albumin (CEA) blocking buffer (0.2 µg/µl CEA, 20 mM Tris HCl pH 7.4-7.8, 150 mM NaCl), rotating for 1 h at 4 °C. Beads were washed with 12 ml cold PBS to remove excess albumin and then incubated with 1 µg biotinylated TREM or an equivalent Flag-C1 medium control. After rotating at 4 °C for 1 h, beads were centrifuged at 1000 g for 1 min, washed in 0.5 ml PBS, and then incubated overnight with 1 ml of the pooled brain lysate rotating at 4 °C. The beads were washed four times with 1 ml 1% NP-40 buffer (centrifuged in between at 1000 g for 1 min), dried with a gauge needle, and resuspended in 50 µl 1% Sodium dodecyl sulfate (SDS) elution buffer (1% SDS, 50 mM Tris pH 8, 5 mM TCEP and 20 mM CAA in MQ). Upon incubation at room temperature for 10 min while shaking and centrifugation at 1000 g for 1 min, 40 µl eluate was aliquoted to a low-binding tube and ramped down to 55 °C for 15 min to alkalize and reduce the proteins. The protein samples were stored at -80 °C until MS analysis. Per TREM protein, 4 biotechnical pulldown replicates were included and handled in a block wise randomized manner.

### Heparin bead pulldowns

Heparin beads (H6508; Sigma-Aldrich) or protein A beads were incubated 5 min at room temperature with 1% NP-40 wash buffer (50mM NaCl, 0.5% NP-40, 5mM MgCl2, 50mM Tris pH 8) and then blocked with 0.5 ml freshly prepared chicken egg albumin (CEA) blocking buffer (0.2 µg/µl CEA, 20 mM Tris HCl pH 7.4-7.8, 150 mM NaCl) rotating for 1 h at 4 °C. Beads were then washed with cold PBS to wash off CEA, and per pulldown, 100 µl of beads was incubated for 1 h at 4 °C with equal amounts of biotinylated TREM ECDs (∼ 200 ng). Beads were washed once with PBS, then washed four times in 1% NP-40 wash buffer, dried with a gauge needle, and resuspended in 1% Sodium dodecyl sulfate (SDS) elution buffer (1% SDS, 50 mM Tris pH 8, 5 mM TCEP and 20 mM CAA in MQ). Supernatants (no bead fraction) were used for western blot analysis.

### Western blot

For validation of TREM-protein concentrations in the medium, beads were dried with a needle, lysed in 5X loading dye (4% SDS, 250 mM Tris HCl pH6.8, 3.33% glycerol (v/v), 10% DTT in deionized water) and boiled for 5 min at 95 °C. Per TREM-medium prep, 15 ul supernatant (no bead fraction) was run on Criterion TGX Stain-Free™ Precast Gels (BIO RAD) in running buffer (1% Tris and glycine (w/v), 1% SDS (v/v) in deionized water), next to 2 μL of Prestained Protein Ladder (PageRuler, Thermo Fisher Scientific) and recombinant biotinylated TREM2 protein (TR2-H82E7-25ug; Acro Biosystems). Samples from the heparin bead immunoprecipitation experiment were prepared similarly. Electrophoresis was performed for 30 min at 90 V and 45 min at 150 V using the PowerPac 1000 (Bio-rad). The proteins were blotted on a midi 13 size low fluorescence polyvinylidene difluoride membrane using Trans-Blot Turbo transfer buffer (BIO-RAD) in the Trans-Blot Turbo Transfer System (BIO RAD), at 2.5 A for 7 min. Blots were incubated with blocking buffer (5% milk powder (w/v) in 0.1% PBS-T) on a shaker for 30 min at room temperature and stained with IRDye® 800CW Streptavidin (1:10,000; Li-COR Biosciences) for 1 h at room temperature. Images were taken using the LI-COR imaging system.

### MS sample preparation

MS sample preparation was performed following the DNA micro spin column suspension trapping (S-Trap) protocol ^35^, with some minor adaptations. Following storage at -80 °C, the TREM - brain lysate IP eluates were incubated at 37 °C for 5 minutes and supplemented with S-Trap lysis buffer up to a final concentration of 5% SDS and 50 mM Tris-HCl pH 8.5. The samples were acidified up to 1.1% phosphoric acid (12% stock solution), mixed with six volumes of binding/washing buffer (90% methanol and 100 mM Tris-HCl pH 8), and loaded onto a plasmid DNA micro column (HiPure from Magen Biotechnology). The protein particulate was retained on the column upon centrifugation at 1400x *g* for 1 minute and the columns were washed four times with binding/washing buffer. Columns were transferred to new LoBind tubes (Eppendorf), supplemented with 0.2 μg Trypsin/Lys-C (Promega) in 50 mM NH4HCO3 and incubated overnight at 37 °C in a humidified incubator. Tryptic peptides were eluted and pooled by subsequent addition of 50 mM NH4HCO3, 0.1% formic acid and 0.1% formic acid in acetonitrile. Collected peptides were dried by SpeedVac and stored at -80 °C.

### LC-MS analysis

Each sample of tryptic digest was redissolved in 0.1% formic acid and 20% was loaded onto an Evotip Pure (Evosep). Peptide samples were separated by standardized 30 samples per day method on the Evosep One liquid chromatography system, using a 15 cm × 150 μm reverse-phase column packed with 1.5 µm C18-beads (EV1137 from Evosep) connected to a 20 µm ID ZDV emitter (Bruker Daltonics). Peptides were electro-sprayed into the timsTOF HT mass spectrometer (Bruker Daltonics) equipped with CaptiveSpray source and measured with the following settings: Scan range 100-1700 m/z, ion mobility 0.65 to 1.5 Vs/cm2, ramp time 100 ms, accumulation time 100 ms, and collision energy decreasing linearly with inverse ion mobility from 59 eV at 1.6 Vs/cm2 to 20 eV at 0.6 Vs/cm2. Operating in dia-PASEF mode, each cycle took 1.38 s and consisted of 1 MS1 full scan and 12 dia-PASEF scans. Each dia-PASEF scan contained two isolation windows, in total covering 300-1200 m/z and ion mobility 0.65 to 1.50 Vs/cm2. Dia-PASEF window placement was optimized using the py-diAID tool ^36^. Ion mobility was auto calibrated at the start of each sample (calibrant m/z, 1/K0: 622.029, 0.992 Vs/cm2; 922.010, 1.199 Vs/cm2; 1221.991, 1.393 Vs/cm2).

### MS data analysis

DIA-PASEF raw data were processed with DIA-NN 1.8.1 ^37, 38^. An in-silico spectral library was generated from the uniprot reference proteome (SwissProt and TrEMBL, canonical and additional isoforms, release 2024-2) using Trypsin/P digestion and at most 1 missed cleavage. Fixed modification was set to carbamidomethylation(C) and variable modifications were oxidation(M) and N-term M excision (at most 1 per peptide). Peptide length was set to 7-30, precursor charge range was set to 2-4, precursor m/z was limited to 280–1220, both MS1 and MS2 mass accuracy were set to 15 ppm, scan window was set to 9, double-pass-mode and match-between-runs (MBR) were enabled. Protein identifiers (isoforms) were used for protein inference. All other settings were left as default.

MS-DAP 1.2 ^39^ was used for downstream analyses of the DIA-NN results. Peptide-level filtering was configured to retain only peptides that were confidently identified in at least 75% of samples within each group of a statistical comparison. Peptide abundance values were normalized using the VSN algorithm, followed by protein-level mode-between normalization. Differential expression analysis (DEA) was performed in a between-subject manner by the MSqRob algorithm ^40^ and resulting p-values were adjusted for multiple testing using the Benjamini–Hochberg False Discovery Rate (FDR) procedure (FDRdea adjusted P-value cutoff of 5%).

All raw peptide detection counts are listed in Supplementary Table 1. To identify TREM-specific interactors, we first filtered for proteins of which the number of measured peptides for the respective TREM-protein was higher than the sum of the number of measured peptides for the other TREMs and control together. Because of the high number of measured interactor proteins for TREML2, an additional filtering was performed here. For this, we selected only proteins of which the number of measured peptides in the TREML2 pulldowns was four times higher than the sum of the number of measured peptides for the other TREMs and control. Interactors were ordered based on their detection confidence score, which was calculated as the percentage of peptides of a specific protein that was identified in all four pulldowns for that respective TREM-protein. As the interactors are expected to be transmembrane or secreted in nature, downstream analyses were performed only for transmembrane and secreted proteins as annotated with Uniprot. Gene Ontology analyses were performed via the online GO platform, and analyses were run for “biological process” and “molecular function”, with False discovery rate correction. STRING analyses were performed via the online platform, with k-means clustering for three clusters and a high confidence criterium (minimum required interaction score: 0.7). Only experimental data was used for STRING analyses. Statistical analysis from the MSqRob analysis and peptide counts are listed in Supplementary Table 1.

## Results

### Identification of novel TREM interactors from human brain lysate

To identify potential receptors and ligands of the TREM-proteins in the brain, we generated recombinant TREM proteins for a pulldown experiment with brain-lysate. Biotinylated extracellular domains (ECD) of the TREM-receptors were produced in HEK293T cells using co-transfection with secreted BirA ligase (Fig 1A), and an empty vector (Flag-C1) was used as a control. The TREMs were successfully secreted and biotinylated, and displayed a complex glycosylation pattern, similar to recombinant TREM2 reference protein (Fig 1B). To identify as many potential interactors as possible, we pooled brain lysates from healthy individuals and Alzheimer’s disease patients from different Braak stages (ranging from 1 to 5) and brain regions (striatum and hippocampus) (Table 1). Four distinct brain tissues (Table 1, Fig 1C) were dounced to extract proteins, and their pooled lysates were incubated with the recombinant TREM proteins. Subsequently, TREM and their interacting proteins were pulled-down using neutravidin beads and analyzed by mass spectrometry (Fig 1D). We identified a range of ∼15000 to ∼35000 peptides per sample (Fig S1A), with clear clustering of the four technical replicates per TREM protein (Fig S1B). After normalization, peptide mapping and annotation, we performed initial filtering of the entire dataset to identify TREM-specific interactors (Fig S1C; Supplementary Table 1). The data from the proteomics yielded interacting proteins that displayed no binding in the control condition, or proteins that showed background binding in the control condition but had increased binding in the TREM condition.

**Figure 1.**
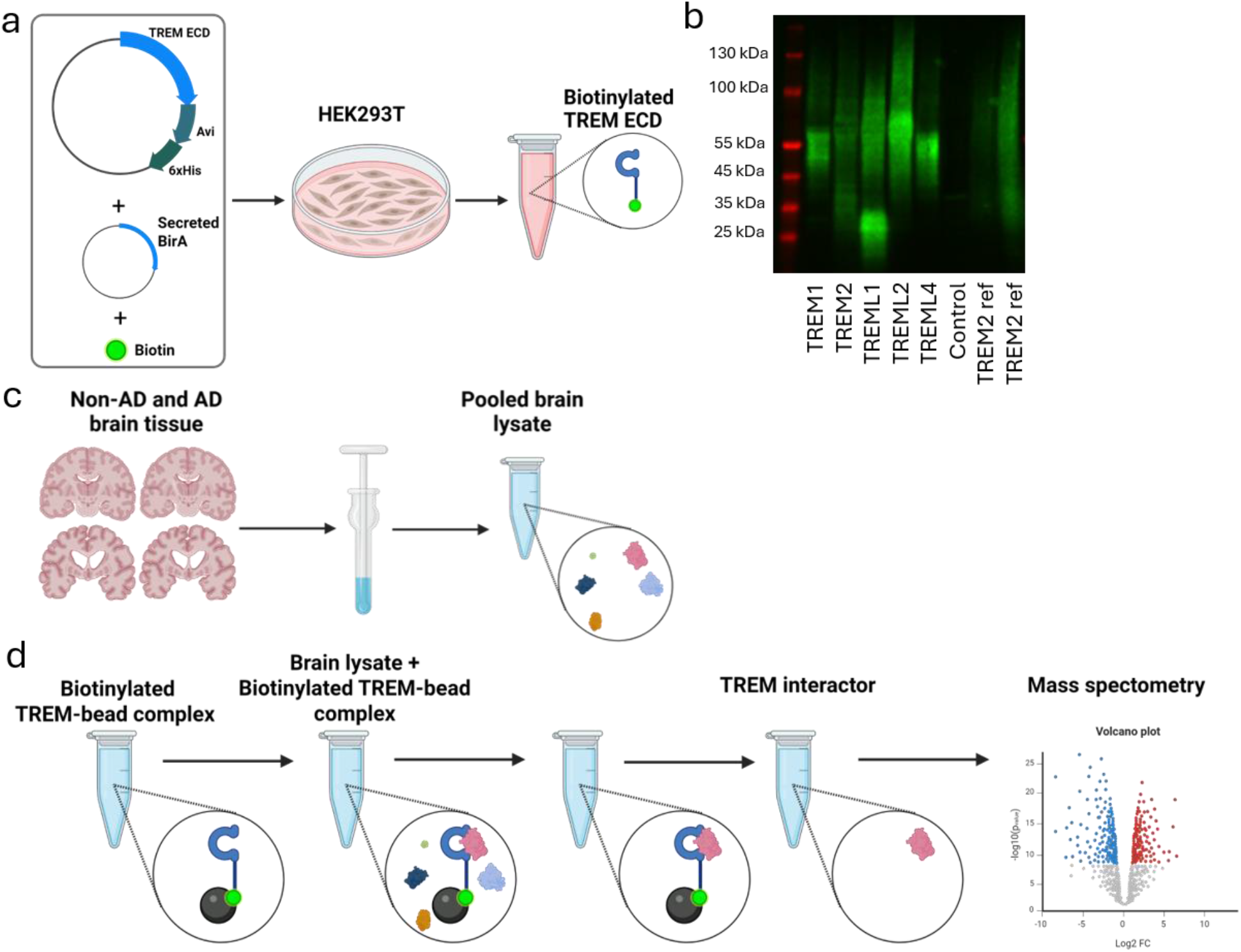
Pipeline for detecting TREM interactors in the brain. A) Biotinylated recombinant TREM proteins were produced in HEK2S3T cells. B) Western blot analysis of biotinylated TREMs in the medium of HEK2S3T cells, detected using Streptavidin aR800. C) AD and non-AD brain tissues were lysed and pooled to generate a wide potential interactor proteome. D) Schematic overview of the pulldown and mass-spectrometry based proteomics workffow.

First, we analyzed the latter by calculating the differential peptide abundance between the control and TREM samples using the MSqRob differential expression algorithm (cut-off log2 fold-change > 2). This required signal in the control conditions, and thus was most suitable for detecting interactors with some background binding. For all TREMs (Fig 2A-E), proteins were identified with significantly increased abundancy in the pulldown compared to the control. For TREML1 the top hits were the three fibrinogen proteins (FGA, FGB, FGG), validating the pulldown approach. Other hits for TREML1 were a set of secreted proteins (FN1, SERPINA1, APOE, A2M), of which several were specific for TREML1 (Fig 2F). Interestingly, the TREM2-interacting apolipoproteins APOE, Clusterin, and APOB interacted with all the TREMs except TREM1, based on either peptide abundance (Fig 2G) or peptide counts (Fig 2H). For TREM2, TREML2, and TREML4, a large group of overlapping proteins was pulled down, which contained mostly intracellular tubulins, keratins, actins and ribosomal proteins, likely reminiscent of non-specific binding.

**Figure 2.**
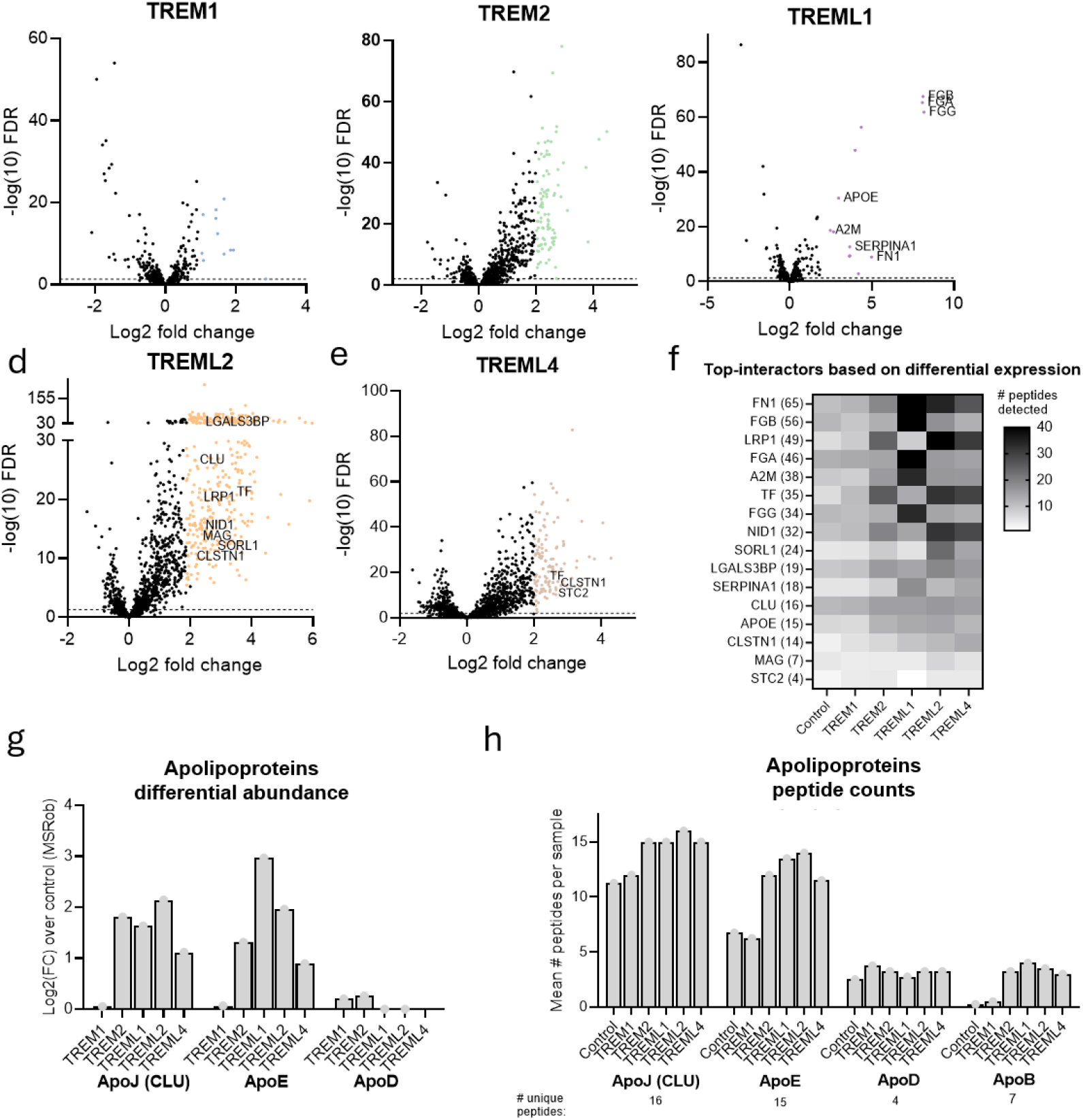
Differential abundance analysis reveals TREM interactors. A-E) Volcano plots depicting differentially abundant proteins compared to the control for the different TREMs. Colored dots represent proteins with a log2 fold-change > 2 (except for TREM1 log2 fold change > 1); transmembrane and secreted proteins are annotated. F) Top-interactors based on Log2 fold change of the number of peptides detected in TREM-pulldown versus control, ranked on the number of unique detected peptides per protein (listed between brackets). Top hits for all TREMs were plotted. G) Log2 fold change for peptide abundance between control and TREMs for APOJ, APOE and ApoD. H) Mean number of detected peptides per sample (averaged over four independent pulldown samples) for APOJ, APOE, ApoD and APOB.

To increase the detection window, we then performed analyses based on the number of peptides identified in the different samples, which allows inclusion of binary interactors in our data (Fig S1C). A specificity score was included to identify TREM-specific binders (interactor peptides having higher counts than all other TREMs and the control combined). In addition, we also selected for transmembrane or secreted proteins (Fig S1C). This selection approach yielded other known interactors for the TREM-proteins, and confirmed PGLYRP1 as a specific interactor for the TREM1 (Fig S2A), and LGALSL and C1Q3L as (specific) interactors for TREM2 (Fig S2B). The known interaction between TREML1 and members of the fibrinogen family was also apparent from these data (Fig S2C). Overall, our pulldown approach identified known interactors for the three most well studied TREMs.

### Interactor analysis for TREM1, TREM2 and TREML4

Next, we searched for new, specific interactors for the different TREM proteins. For TREM1, TREM2 and TREML4, the number of confidently detected specific interactor proteins after data filtering (Fig S1C) was very low. For TREM1, only TREM1 itself and PGLYRP1 specifically precipitated, at least verifying the TREM1/PGLYRP1 interaction in the human brain. For TREM2, Nucleobindin 2 (NUCB2) appeared as a possible secreted interactor (Fig 3B). However, its specificity over TREML2 and TREML4 was low, so it is not likely to be an exclusive interactor. Also for TREML4 no specific interactor was identified. ITM2B, a microglial transmembrane protein with high expression on microglia, displayed some specificity, but also here some background was detected in the TREM2 and TREML2 conditions. Additional TREML4-binding partners were identified based on lower unique peptides counts, conveying even less confident binders (Fig 3C). Whereas the number of strong TREM1, TREM2 and TREML4 interactors is sparse, the identification of the respective TREM bait protein in each interactor dataset confirms the accuracy of the experimental set-up.

**Figure 3.**
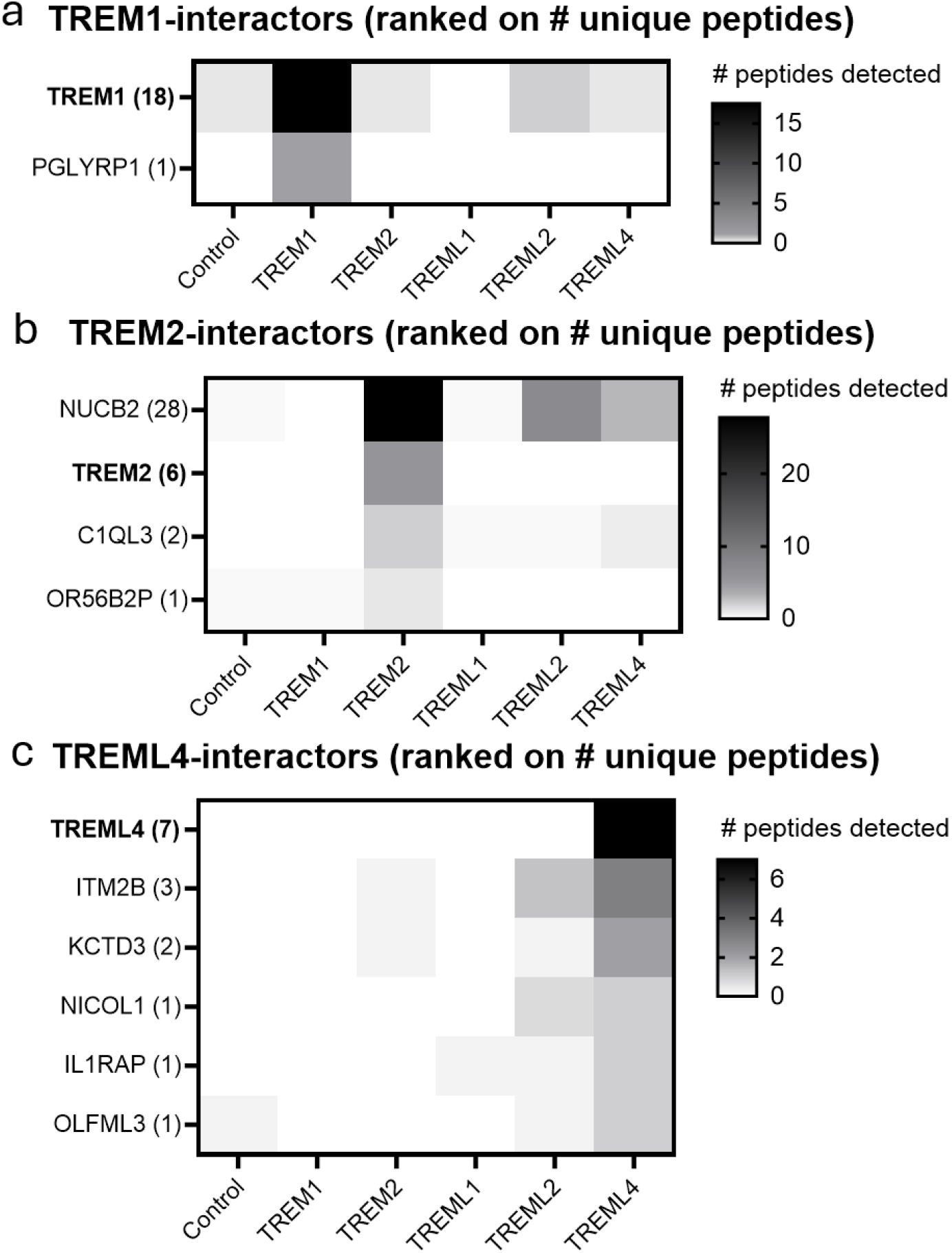
Detection of specific interactors for TREM1, TREM2 and TREML4. A-C) Specific interactors for TREM1, TREM2 orTREML4 based on a peptide detection score of > S0% among replicates, ranked on number of unique peptides (listed between brackets after each protein).

### Specific interactions of TREML1 with lipoproteins and complement proteins

For TREML1, a multitude of very specific interactors was identified (Fig 4A). Gene ontology analysis on these hits identified complement signaling as a common molecular function of a large set of these proteins, with nearly all members of the classical complement pathway identified (CFH, CP, CPS1, C9, C7, CFB, C2, C4BPA, 8B, F9, PON1, C1R) (Fig 4B).

**Figure 4.**
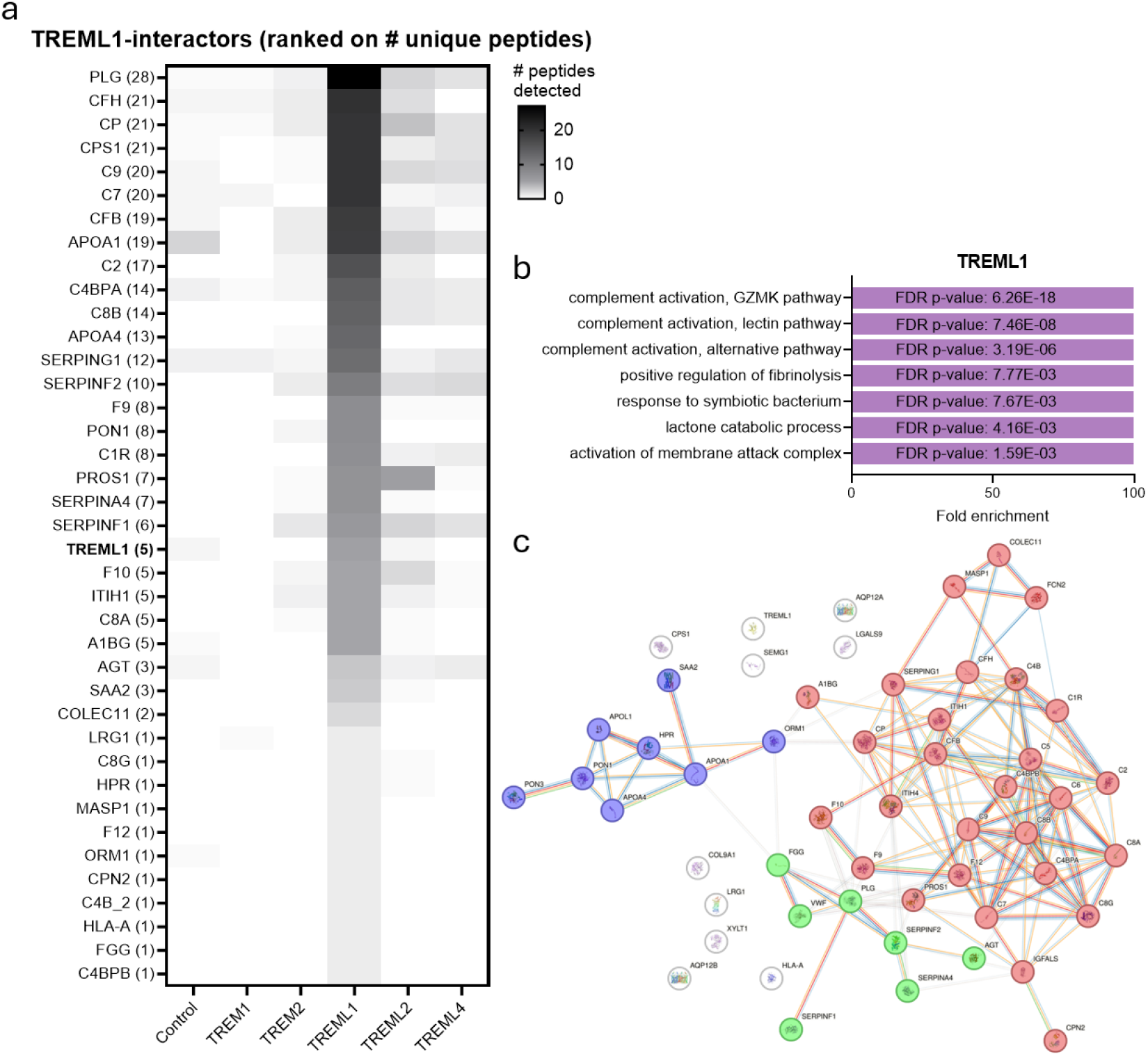
TREML1 binds to complement proteins and apolipoproteins. A) TREML1 top-interactors with a peptide detection score of > S0%, ranked on number of unique peptides (listed between brackets). B) Fold enrichment scores and FDR-corrected significance for TREML1 Gene Ontology analysis pathways, based on the entire filtered set (25-100% confidence). C) STRING analysis for top-interactors depicted in panel A, k-means clustered into three groups.

In addition, several TREML1 interactors were linked to blood clotting, as indicated by the GO pathway “positive regulation of fibrinolysis” (Fig 4B), and probably precipitated together with the fibrinogens. Lastly, a set of apolipoproteins (APOA1 and APOA4) – components of high density lipoprotein particles - was identified as specific TREML1 interactors (Fig 4B). STRING analysis with k-means clustering (PPI enrichment p-value:< 1.0e-64) showed that the fibrinogen family, the complement system and the APOA1 and APOA4 proteins are part of a larger interaction network (Fig 3D).

### TREML2 is a novel interactor of heparan sulfate proteoglycans

Also for TREML2 a large number of specific interactors was identified. To increase the likelihood for a direct interaction, a more stringent selectivity score was used (Fig S1C), resulting in 22 interactors (Fig 5A). Strikingly, these interactors included many different Glypicans (GPC3, GPC4, GPC6), Syndecans (SDC2, SDC3, SDC4), Neurexins (NRXN1, NRXN2, NRXN3), and SPOCK proteins (SPOCK1, SPOCK2, SPOCK3), which are all proteins modified with heparan sulfates. A GO analysis revealed pathways surrounding these hits, with primarily “Neuroligin family protein binding” (NRXN1, NRXN2, NRXN3) and “Metalloendopeptidase activity” (SPOCK1, SPOCK2, SPOCK3) enriched (Fig 5B). Other striking hits were the TREML2-specific binding to the lipid-receptors LRP2 and LDLR (Fig 5A). Even though these neuroligins, metalloendopeptidases, lipid receptors, glypicans and syndecans do not necessarily bind amongst each other (Fig 5C; PPI enrichment p-value STRING analysis: 1.61e-27), they all share the important feature that they are all heparan sulfate proteoglycans (HSPGs), or form a complex with them.

**Figure 5.**
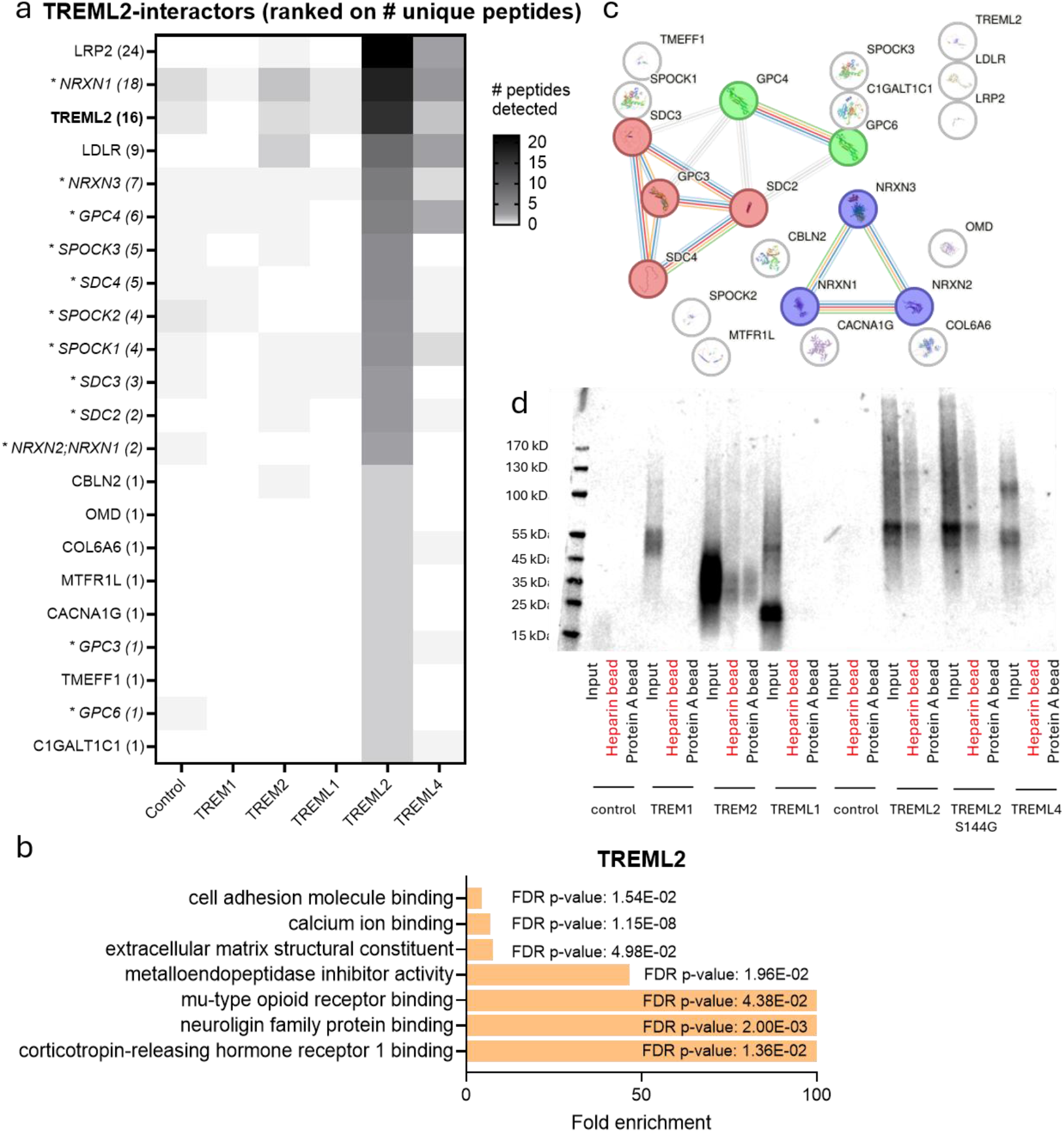
TREML2 interacts with heparin and heparan sulfate proteoglycans. A) TREML2 top-interactors with a peptide detection score of > S0% and stringent filtering for specificity. Hits are ranked on number of unique peptides (listed between brackets). Heparan sulfate modified proteins (HSPGs) are depicted in italic and with an asterisk. B) Fold enrichment scores and FDR-corrected significance for TREML2 Gene Ontology analysis pathways, based on the entire filtered set (25-100% confidence). C) STRING analysis for top-interactors depicted in panel A, k-means clustered into three groups. D) Pulldown of the TREM proteins with heparin beads and protein A beads, to detect specific interactions with heparan sulfates.

Therefore, we hypothesized that TREML2 may bind to heparan sulfate chains on all of these proteins. To test this, the ability of the different TREM proteins to interact with heparin beads was assessed. Indeed, while TREM1, TREML1 and TREML4 did not interact with heparin beads, TREML2 displayed robust and specific binding to the heparin beads (Fig 4D). TREM2 interacted with both the Prot A and the heparin beads, disallowing any statements about its ability to interact. To test whether the protective variant S144G affects the binding to heparin, we tested TREML2 S144G alongside the wild-type protein. No difference in binding was observed between both variants, suggesting that the affinity of the variant TREML2 is not majorly affected. Together, these data show that TREML2 interacts with a multitude of HSPGs, probably via its ability to bind to their heparan sulfate moieties.

## Discussion

The TREM receptor family plays a key role in the activation and modulation of myeloid cells, and especially TREM1, TREM2 and TREML2 have been linked to neuroinflammation and AD. By interacting with different ligands, ranging from aggregated proteins to pattern recognition molecules, they determine the inflammatory environment of the brain. Here, using interaction proteomics with healthy and AD-diseased human brain samples, we confirmed several known interactors of the different TREMs, and identified novel interactors for TREML1 and TREML2. TREML1 pulled down the apolipoproteins APOA1 and APOA4, as well as several components of the complement system. TREML2, on the other hand, interacted with a multitude of proteins modified with heparan sulfate chains, and was confirmed to directly associate with heparin. In contrast to APOE and other TREM2 ligands, we did not identify an interaction with the potential TREM2 receptor Transgelin-2^41^.

A key function of the most well-studied member of the TREM family, TREM2, is its sensing and uptake of lipoproteins ^42^. Our data show that besides TREM2, also the TREML-proteins, but not TREM1, can interact with APOE, APOB, and APOJ, arguing for a broader role of the TREMs in apolipoprotein biology. Interestingly, the lipoproteins APOA1 and APOA4 specifically interacted with TREML1, with very high peptide scores. While TREML1 and APOA1/APOA4 are expressed in the brain or can be imported over the blood-brain barrier, this interaction is more likely to be relevant in the periphery. TREML1 is predominantly expressed on blood platelets and increases blood clotting via its known fibrinogen interactions, while APOA1 and APOA4 have both been shown to inhibit platelet activation and blood clotting ^43–45^. It is possible that APOA1 and APOA4 bind TREML1 competitively with fibrinogens, hereby preventing platelet aggregation. Alternatively, TREML1 interacts with APOA1/4 indirectly, for example via αIIbβ3 Integrin that is expressed on platelets and demonstrated to interact with APOA4^46^. However, no integrins were identified in our TREML1 pulldowns, suggesting an indirect interaction is probably not modulated via integrin interactions.

Aside from the apolipoproteins, we confirmed that TREML1 in brain lysate pulls down the known interaction partners the fibrinogens (FGA, FGB and FGG), and several other regulators of blood coagulation, such as fibrin (FN1), Plasminogen (PLG) and Serpins. Whether TREML1 acts on multiple stages of the coagulation pathway is unknown, but it seems plausible that TREML1 interacts with the other coagulation members via the fibrinogens. Similarly, TREML1 interacted with nearly all members of the classical complement system, a system known to interact with fibrinogen ^47^. Also this interaction was highly specific for TREML1, and argues for a broader role for TREML1 in innate immunity in the periphery and brain, since the complement system plays an important role in synaptic pruning and brain homeostasis ^48^. Given the presence of blood vessels in the brain, it is assumed these interactions primarily occur in the vessels, but they could also occur in the brain upon rupture of the blood-brain barrier.

For TREML2, the heparan sulfate proteoglycans we identified as interactors mediate synaptic function (NRXN neuroligin receptors), lipid transport (LDLR, LRP4, LRP1), cellular growth pathways (Glypicans), and extracellular matrix and focal adhesion (Syndecans). Tying the pulldown of these interactors together, we demonstrate that TREML2 interacts with heparin, making it likely that TREML2 interacts with the heparan sulfate chains on these proteins. None of the other TREMs recapitulated this interaction, in both the proteomics and our *in vitro* validation with heparin pulldowns, arguing for a specific role of TREML2 in these interactions. Due to the strong negative charge of heparan sulfate chains, interactions of proteins with HSPGs are usually orchestrated by basic arginine and lysine residues from the interacting protein side. In line with this, out of all the TREMs, the ectodomain of TREML2 has the highest isoelectic point (pI) at 9.9 ^49^, indicating it has the most positive charge at neutral pH with a surplus of 15 basic residues. Functionally, TREML2 and HSPGs could be linked to regulate AD risk, as the HS3ST1 gene involved in generating 3-O-sulfated HSPGs is genetically linked to AD risk ^50, 51^. HSPGs are also known to interact with Tau and APOE, and are involved in the aggregation of amyloid-β ^52–54^. Thus, TREML2 could potentially influence some of these interactions. However, while the protective S144G TREML2 variant is situated in the ligand-binding domain, it did not severely affect heparin binding in our pulldown assays. Direct affinity measurements should be used to resolve whether small, but possibly relevant, affinity differences between the wildtype and S144G TREML2 variants are present. While the interactions between TREML2 and HSPGs seem robust, we cannot exclude that TREML2 has specific ligands that drive its activation. Our pulldowns did not identify any directly relevant, specific interactors, but given the large interactor list and the notion that TREML2 interacted with different lipoproteins it seems likely that it has other relevant binding partners. Together, our pulldowns identified an interaction of TREML2 with HSPGs, which could be of relevance in the development of AD. Further experiments defining the interactions in more detail will be necessary to determine this.

The TREM-receptor family plays a crucial role in fine-tuning inflammation and homeostasis in the brain, either directly on microglia, or via peripheral immune cell influx upon a compromised the blood-brain-barrier. Here, using affinity purification proteomics we confirmed the interaction of TREM1 with PGLYRP1 in human brain lysates, and identified novel interactions for TREML1 and TREML2. TREML2, which is genetically linked to AD, was found to interact with heparan sulfates, which are also genetically linked to AD, potentially linking both AD risk genes in a shared process.

### Limitations

While our approach used recombinant TREM proteins produced in human cells, it could be that the cell types expressing the TREM-receptors endogenously produce a different TREM-glycosylation pattern, allowing interactions to go unnoticed. Furthermore, low-affinity interactions, or detergent sensitive interactions, are likely not sustained during our protein incubation or washes. Alternatively, the TREM proteins could interact with ligands that are of a different nature, like lipids or metabolites. A similar approach using other types of -omics analyses could identify such interactions. Lastly, we are unsure whether the multitude of common interactors between TREM2, TREML2, and TREML4 has biological relevance or was inherent to the current experimental set-up.

## Acknowledgements

We thank the Netherlands brain bank for providing the tissue blocks and we thank the donors for the contribution of their brain tissue to science. We thank Aniko Mikes for helping set-up the heparin-bead immunoprecipitations, and members of the Candidate Center lab and Rik van der Kant lab for helpful discussions. This work was supported by an Alzheimer Nederland pilot grant for R.H.W. and a ZonMw Dementia Fellowship for K.M.A.K.

---

**Figure S1.**
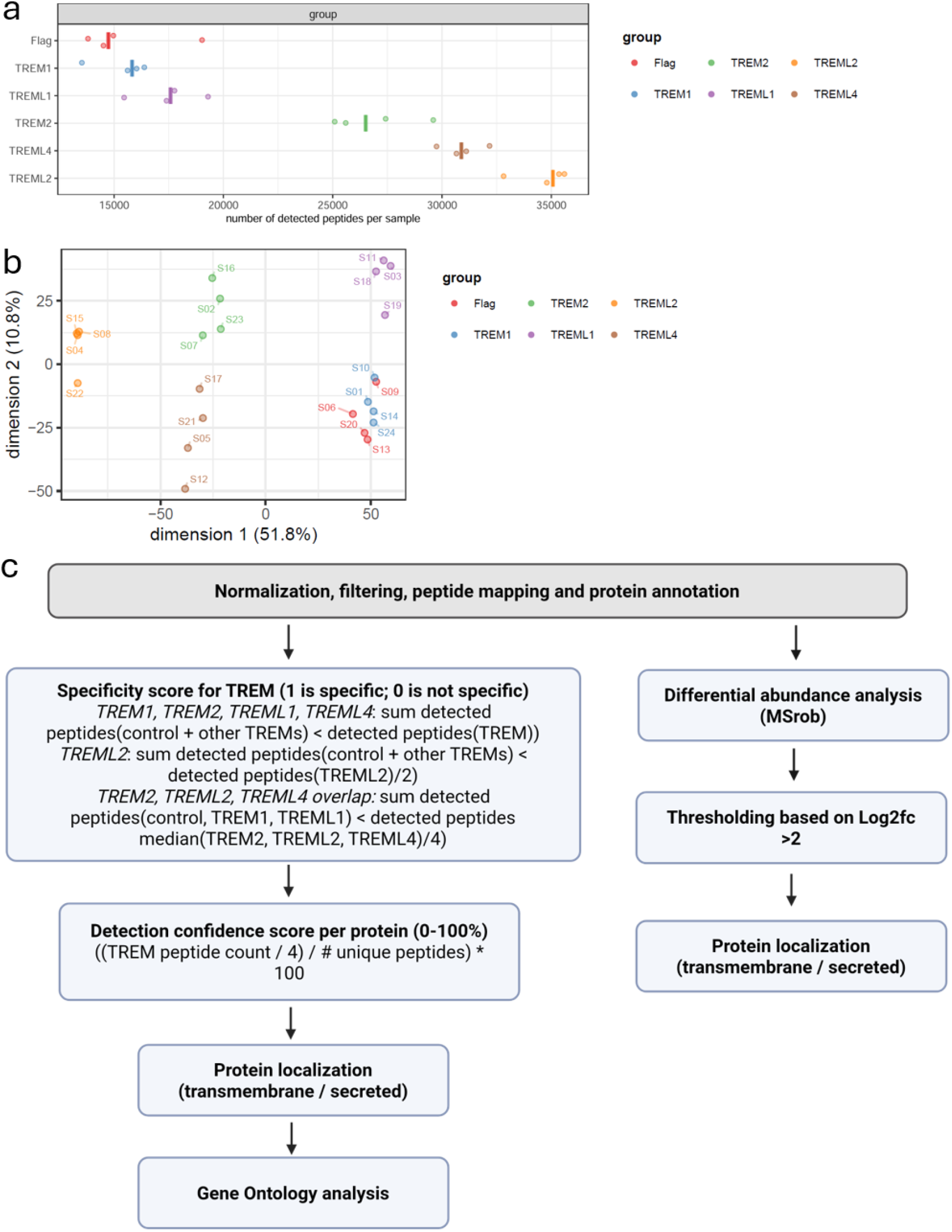
Proteomics quality control and analysis pipeline. A) Number of detected peptides per sample. B) Principial component analysis reffecting separation of TREM and control pulldown datasets based on the number of identified peptides per sample. C) Schematic of the two data analysis approaches after data pre-processing. The left pipeline was designed to primarily detect specific interactions, where the number of identified peptides for a specific TREM had to be higher than the sum of all other TREMs and the control combined. The right analysis was used to detect binders with increased binding of a TREM compared to the control, but did not include proteins not detected in the control samples (binary interactions).

**Figure S2.**
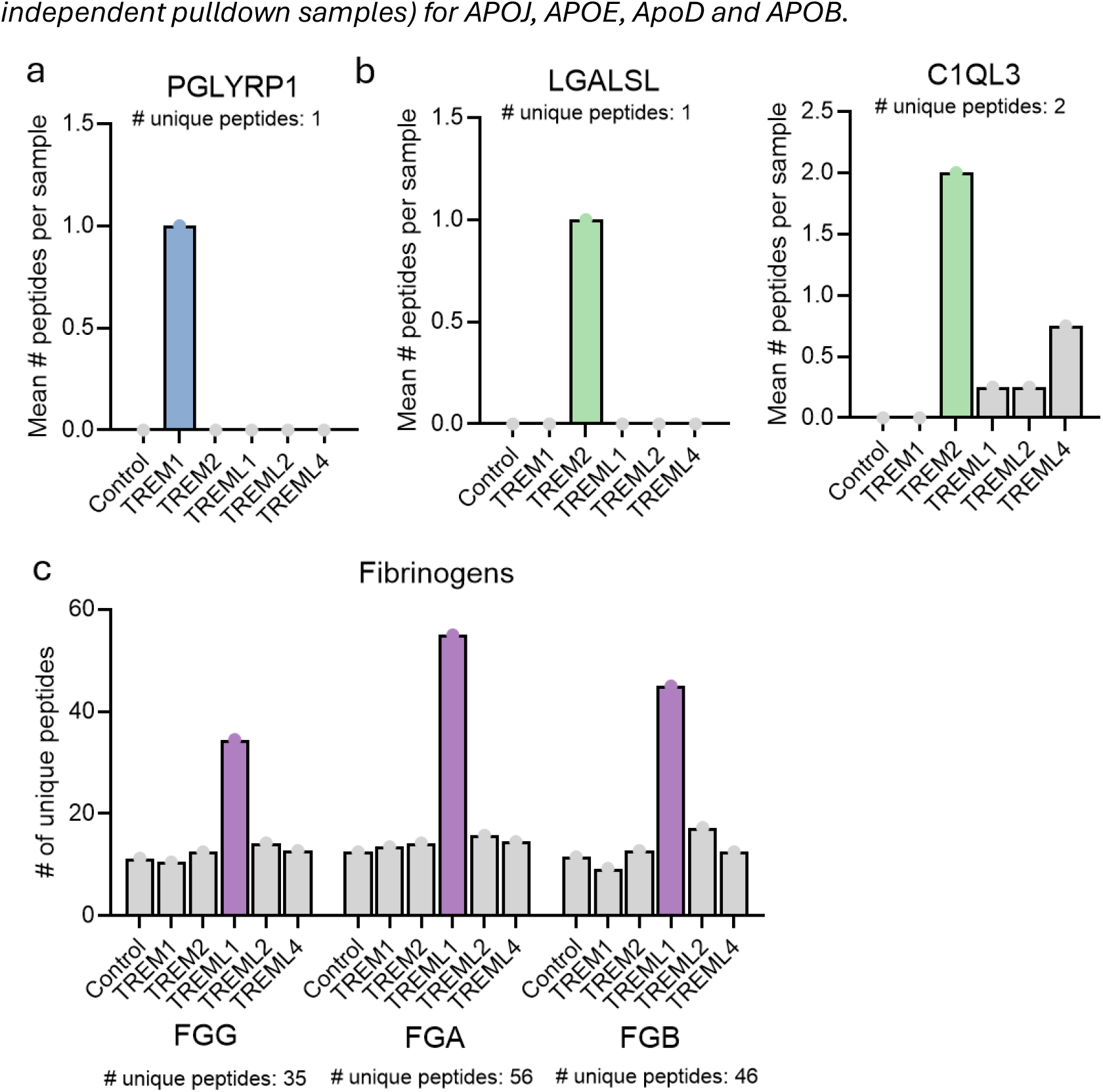
Identification of known TREM protein interactors. Mean number of detected peptides per sample (averaged over four independent pulldown samples) for A) PGLYRP1, B) LGALSL and C1QL3, and C) FGG, FGA and FGB (fibrinogen protein family members).

## References

1. Colonna, M. The biology of TREM receptors. Nat Rev Immunol 23, 580–594 (2023).

2. Klesney-Tait, J., Turnbull, I.R. & Colonna, M. The TREM receptor family and signal integration. Nat Immunol 7, 1266–1273 (2006).

3. Hemmi, H. et al. A New Triggering Receptor Expressed on Myeloid Cells (Trem) Family Member, Trem-Like 4, Binds to Dead Cells and Is a DNAX Activation Protein 12-Linked Marker for Subsets of Mouse Macrophages and Dendritic Cells. The Journal of Immunology 182, 1278–1286 (2009).

4. Bouchon, A., Dietrich, J. & Colonna, M. Cutting edge: inflammatory responses can be triggered by TREM-1, a novel receptor expressed on neutrophils and monocytes. J Immunol 164, 4991–4995 (2000).

5. Daws, M.R., Lanier, L.L., Seaman, W.E. & Ryan, J.C. Cloning and characterization of a novel mouse myeloid DAP12-associated receptor family. Eur J Immunol 31, 783–791 (2001).

6. Wilson, E.N. et al. TREM1 disrupts myeloid bioenergetics and cognitive function in aging and Alzheimer disease mouse models. Nat Neurosci 27, 873–885 (2024).

7. Jonsson, T. et al. Variant of TREM2 associated with the risk of Alzheimer’s disease. N Engl J Med 368, 107–116 (2013).

8. Del Campo, M. et al. CSF proteome profiling across the Alzheimer’s disease spectrum reflects the multifactorial nature of the disease and identifies specific biomarker panels. Nat Aging 2, 1040–1053 (2022).

9. Feiten, A.F. et al. TREM2 expression level is critical for microglial state, metabolic capacity and efficacy of TREM2 agonism. Nature Communications 17, 2002 (2026).

10. Keren-Shaul, H. et al. A Unique Microglia Type Associated with Restricting Development of AlzheimerC#x2019;s Disease. Cell 169, 1276–1290.e1217 (2017).

11. Guerreiro, R. et al. TREM2 variants in Alzheimer’s disease. N Engl J Med 368, 117–127 (2013).

12. Sims, R. et al. Rare coding variants in PLCG2, ABI3, and TREM2 implicate microglial-mediated innate immunity in Alzheimer’s disease. Nat Genet 49, 1373–1384 (2017).

13. Wang, Y. et al. TREM2 lipid sensing sustains the microglial response in an Alzheimer’s disease model. Cell 160, 1061–1071 (2015).

14. Zhong, L. et al. TREM2 receptor protects against complement-mediated synaptic loss by binding to complement C1q during neurodegeneration. Immunity 56, 1794–1808.e1798 (2023).

15. Boza-Serrano, A. et al. Galectin-3, a novel endogenous TREM2 ligand, detrimentally regulates inflammatory response in Alzheimer’s disease. Acta Neuropathol 138, 251–273 (2019).

16. Atagi, Y. et al. Apolipoprotein E Is a Ligand for Triggering Receptor Expressed on Myeloid Cells 2 (TREM2). J Biol Chem 290, 26043–26050 (2015).

17. Yeh, F.L., Wang, Y., Tom, I., Gonzalez, L.C. & Sheng, M. TREM2 Binds to Apolipoproteins, Including APOE and CLU/APOJ, and Thereby Facilitates Uptake of Amyloid-Beta by Microglia. Neuron **91**, 328–340 (2016).

18. Filipello, F. et al. The Microglial Innate Immune Receptor TREM2 Is Required for Synapse Elimination and Normal Brain Connectivity. Immunity 48, 979–991.e978 (2018).

19. Bhusal, A. et al. The microglial innate immune protein PGLYRP1 mediates neuroinflammation and consequent behavioral changes. Cell Rep 43, 113813 (2024).

20. Washington, A.V. et al. TREM-like transcript-1 protects against inflammation-associated hemorrhage by facilitating platelet aggregation in mice and humans. J Clin Invest **119**, 1489–1501 (2009).

21. Derive, M. et al. Soluble TREM-like transcript-1 regulates leukocyte activation and controls microbial sepsis. J Immunol 188, 5585–5592 (2012).

22. Carrasquillo, M.M. et al. A candidate regulatory variant at the TREM gene cluster associates with decreased Alzheimer’s disease risk and increased TREML1 and TREM2 brain gene expression. Alzheimers Dement 13, 663–673 (2017).

23. Xu, Z.H. et al. TREML2 Modulates Microglial Phagocytosis of Myelin Debris via TLR9 in Chronic Cerebral Ischemia. Inflammation **49**, 4 (2025).

24. Fu, X.X. et al. Knockdown of TREML2 Alleviates Neuropathological Hallmarks and Cognitive Deficiency in a Model of Sporadic Alzheimer’s Disease. J Inflamm Res 17, 10471–10478 (2024).

25. Jiang, T. et al. A Missense Variant in TREML2 Reduces Risk of Alzheimer’s Disease in a Han Chinese Population. Mol Neurobiol 54, 977–982 (2017).

26. Li, J.Q. et al. Associations Between TREML2 Gene Variants and Alzheimer’s Disease: Biomarkers, Neuroimage, and Cognition. J Alzheimers Dis **96**, 1555–1563 (2023).

27. Nedeva, C. et al. TREML4 receptor regulates inflammation and innate immune cell death during polymicrobial sepsis. Nat Immunol 21, 1585–1596 (2020).

28. Ramirez-Ortiz, Z.G. et al. The receptor TREML4 amplifies TLR7-mediated signaling during antiviral responses and autoimmunity. Nat Immunol 16, 495–504 (2015).

29. Schlepckow, K. et al. An Alzheimer-associated TREM2 variant occurs at the ADAM cleavage site and affects shedding and phagocytic function. EMBO Mol Med **9**, 1356-1365 (2017).

30. Ewers, M. et al. Increased soluble TREM2 in cerebrospinal fluid is associated with reduced cognitive and clinical decline in Alzheimer’s disease. Sci Transl Med 11 (2019).

31. Zhong, L. et al. Soluble TREM2 induces inflammatory responses and enhances microglial survival. J Exp Med 214, 597–607 (2017).

32. Zhong, L. et al. Soluble TREM2 ameliorates pathological phenotypes by modulating microglial functions in an Alzheimer’s disease model. Nat Commun 10, 1365 (2019).

33. Lu, L., Liu, X., Fu, J., Liang, J., Hou, Y. & Dou, H. sTREM-1 promotes the phagocytic function of microglia to induce hippocampus damage via the PI3K-AKT signaling pathway. Sci Rep 12, 7047 (2022).

34. Wijdeven, R.H. et al. Cholesterol and ORP1L-mediated ER contact sites control autophagosome transport and fusion with the endocytic pathway. Nat Commun 7, 11808 (2016).

35. Thanou, E. et al. Suspension TRAPping Filter (sTRAP) Sample Preparation for Quantitative Proteomics in the Low µg Input Range Using a Plasmid DNA Micro-Spin Column: Analysis of the Hippocampus from the 5xFAD Alzheimer’s Disease Mouse Model. Cells 12 (2023).

36. Skowronek, P. et al. Rapid and In-Depth Coverage of the (Phospho-)Proteome With Deep Libraries and Optimal Window Design for dia-PASEF. Mol Cell Proteomics 21, 100279 (2022).

37. Demichev, V. et al. dia-PASEF data analysis using FragPipe and DIA-NN for deep proteomics of low sample amounts. Nat Commun 13, 3944 (2022).

38. Demichev, V., Messner, C.B., Vernardis, S.I., Lilley, K.S. & Ralser, M. DIA-NN: neural networks and interference correction enable deep proteome coverage in high throughput. Nat Methods 17, 41–44 (2020).

39. Koopmans, F., Li, K.W., Klaassen, R.V. & Smit, A.B. MS-DAP Platform for Downstream Data Analysis of Label-Free Proteomics Uncovers Optimal Workflows in Benchmark Data Sets and Increased Sensitivity in Analysis of Alzheimer’s Biomarker Data. J Proteome Res 22, 374–386 (2023).

40. Goeminne, L.J.E., Sticker, A., Martens, L., Gevaert, K. & Clement, L. MSqRob Takes the Missing Hurdle: Uniting Intensity- and Count-Based Proteomics. Anal Chem **92**, 6278–6287 (2020).

41. Zhang, X. et al. Soluble TREM2 ameliorates tau phosphorylation and cognitive deficits through activating transgelin-2 in Alzheimer’s disease. Nature Communications 14, 6670 (2023).

42. Abdulkhaliq, A.A. et al. TREM2 in neurodegeneration and diseases. Molecular Psychiatry 31, 3978–3986 (2026).

43. Jones, W.L. et al. Apolipoprotein A-I, elevated in trauma patients, inhibits platelet activation and decreases clot strength. Platelets 33, 1119–1131 (2022).

44. Xu, X.R. et al. Apolipoprotein A-IV binds αIIbβ3 integrin and inhibits thrombosis. Nature Communications **9**, 3608 (2018).

45. Yan, M. et al. Apolipoprotein A-IV regulates coagulation and ischemic stroke by potentiating activated protein C. Journal of Thrombosis and Haemostasis 23, 2837–2848 (2025).

46. Xu, X.R. et al. Apolipoprotein A-IV binds αIIbβ3 integrin and inhibits thrombosis. Nat Commun **9**, 3608 (2018).

47. King, R.J. et al. Fibrinogen interaction with complement C3: a potential therapeutic target to reduce thrombosis risk. Haematologica 106, 1616–1623 (2021).

48. Stevens, B. et al. The Classical Complement Cascade Mediates CNS Synapse Elimination. Cell 131, 1164–1178 (2007).

49. Kober, D.L. et al. Neurodegenerative disease mutations in TREM2 reveal a functional surface and distinct loss-of-function mechanisms. eLife 5, e20391 (2016).

50. Wang, Z. et al. Increased 3-O-sulfated heparan sulfate in Alzheimer’s disease brain is associated with genetic risk gene HS3ST1. *Sci Adv* **9**, eadf6232 (2023).

51. Readhead, B. et al. Heparin treatment is associated with a delayed diagnosis of Alzheimer’s dementia in electronic health records from two large United States health systems. Molecular Psychiatry 30, 1461-1465 (2025).

52. Ozsan McMillan, I., Li, J.P. & Wang, L. Heparan sulfate proteoglycan in Alzheimer’s disease: aberrant expression and functions in molecular pathways related to amyloid-β metabolism. Am J Physiol Cell Physiol 324, C893–c909 (2023).

53. Libeu, C.P. et al. New Insights into the Heparan Sulfate Proteoglycan-binding Activity of Apolipoprotein E*. Journal of Biological Chemistry 276, 39138–39144 (2001).

54. Zhao, J. et al. 3-O-Sulfation of Heparan Sulfate Enhances Tau Interaction and Cellular Uptake. Angew Chem Int Ed Engl **59**, 1818–1827 (2020).

